# COPAL: An ensemble of protein-ligand co-folding models enriches preferred ligand predictions for the LuxR-family of quorum sensing receptors

**DOI:** 10.64898/2026.09.04.749498

**Authors:** Davi Nakajima An, Amy L. Schaefer, Szu-Min Chang, Bianxia Bai, Charlotte I. Mulligan, Yasuhiro Oda, E. Peter Greenberg, Frank DiMaio

## Abstract

Acyl-homoserine lactone (AHL) quorum sensing enables many species of proteobacteria to coordinate collective behaviors. In such systems, a synthase produces an AHL signal, which is sensed by a LuxR-family receptor. Despite extensive genomic annotation of LuxR homologs, preferred AHLs for most receptors remain unknown, limiting functional understanding of quorum sensing across diverse bacteria. Here, we present the COPAL (combining ordered predictions of audited ligands) pipeline, which integrates multiple protein-ligand co-folding models to identify preferred AHLs for a specific LuxR. Benchmarking on a leakage-controlled subset of 96 experimentally characterized LuxR-AHL pairs shows that COPAL places the preferred AHL within the top-6 candidates (out of 58) for 68% of receptors, outperforming every individual co-folding model. Further, inter-model agreement correlates with ranking accuracy, offering an indication of confidence. We show that COPAL resolves the specificity shift induced by three-point mutations in LasR and correctly nominates C8-HSL as the preferred ligand for the previously uncharacterized *Mesorhizobium* sp. NJ3 receptor, which we verified experimentally. Finally, we release the Ranked AHL-LuxR Prediction Hub (RALPH), comprising precomputed rankings for about 10,000 unique LuxR homologs. More broadly, COPAL shows that unweighted rank aggregation of complementary co-folding models offers a general strategy for predicting receptor-ligand specificity in data-scarce biological systems.

## Introduction

Predicting preferred ligand-receptor interactions among signaling proteins is a long-standing problem in biology, with applications spanning therapeutic target discovery to modeling native receptor-ligand interactions in the cell^1^. Within this set of problems lies the ligand-receptor interactions in the bacterial communication systems known as LuxR-LuxI-type quorum sensing (QS)^2–4^. In such QS systems (overview in Figure 1a), an acyl-homoserine lactone (AHL) signal is synthesized by a LuxI-type enzyme and freely diffuses out of and into cells at a sufficient concentration the AHL can bind to a cognate LuxR-family receptor. The LuxR-type receptors are transcriptional regulators controlling a variety of downstream behaviors including conjugation, exoenzyme production, or antibiotic synthesis. Specificity of a given QS system is a function of the AHL acyl group moiety (Figure 1b), which varies both in length (4 to 20 carbons) and chemical substituents, including aromatic and branched-chain groups^5,6^.

**Figure 1.**
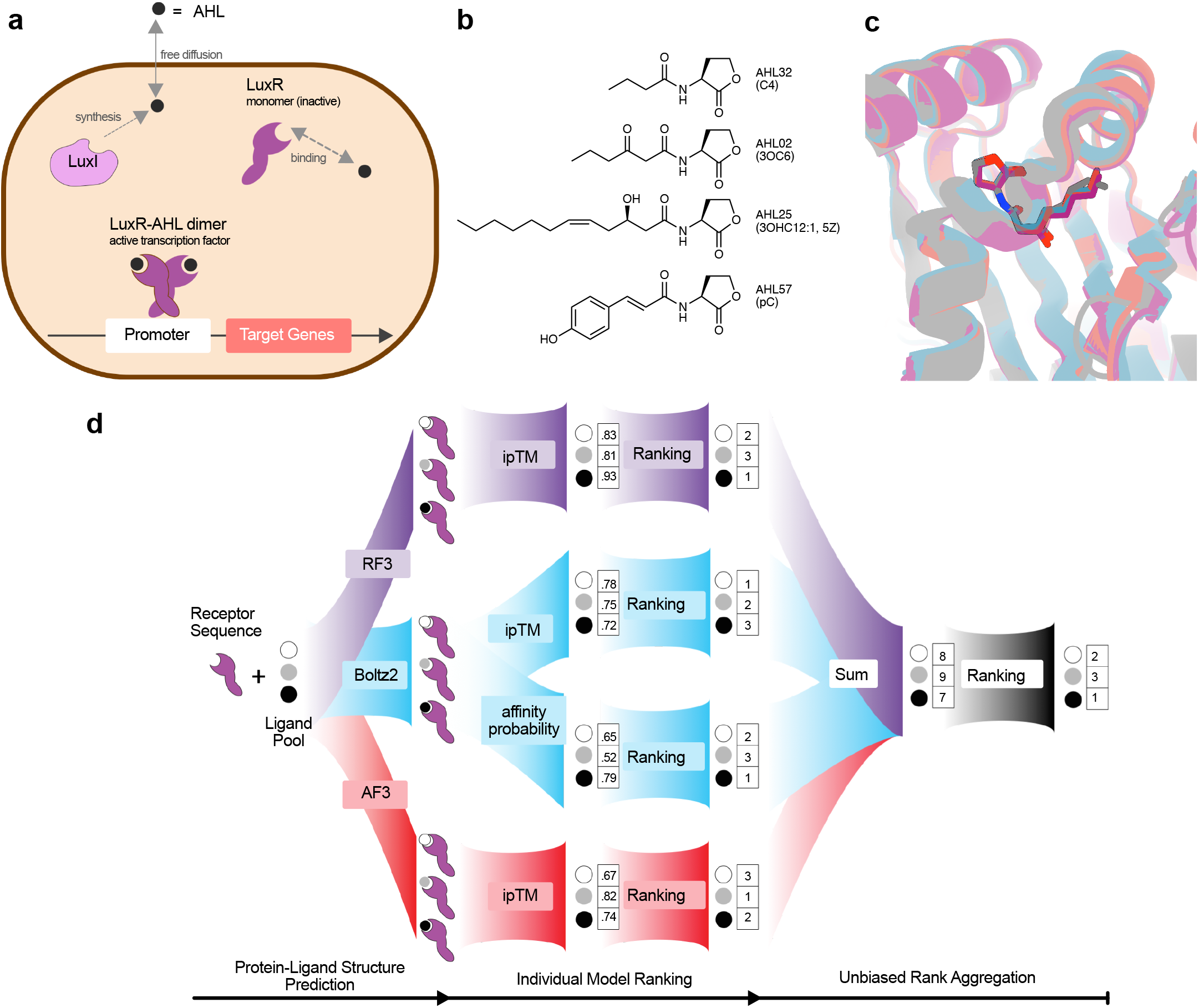
Overview of AHL-type quorum sensing and the COPAL pipeline. **a)** Generalized model of quorum sensing: LuxI synthesizes acyl-homoserine lactone (AHL) signals that diffuse across the cell membrane, accumulate to a threshold concentration and then bind the cytoplasmic receptor LuxR. The LuxR-AHL complex then regulates target gene transcription. **b)** Chemical structures of four representative AHLs in the ligand pool (complete list in Supplemental Table S1) **c)** Model predictions (RF3 in purple, Boltz2 in blue, AF3 in red) of Agro_001 with its AHL_pref_ 3OC8 superimposed with the crystal structure (in gray) from PDBid: 1L3L **d)** COPAL pipeline schematic. A protein sequence is input with a curated pool of candidate ligands to three independent structure prediction platforms (RoseTTAFold3, Boltz2, AlphaFold3). Each platform generates protein-ligand complex models from which binding confidence metrics are extracted: ipTM scores for RF3 and AF3, and both ipTM scores and binding probability predictions for Boltz2. These metrics are used to rank the ligand pool independently within each model, producing four individual ranked lists. Rankings are then aggregated by unweighted rank summation to yield a final COPAL ranking.

Determining which AHL a given LuxR receptor recognizes is important for understanding these QS systems, but it is a challenging problem for current computational methods. LuxR proteins share conserved ligand-binding pockets (e.g. the structure shown in Figure 1c; gray) that interact with structurally similar AHLs^7,8^, yet highly similar LuxRs can bind distinct AHLs and homologs sharing a single AHL can show as low as 25% sequence identity^8,9^. Compounding this, orphan LuxRs lacking a nearby *luxI* gene and receptors that bind multiple AHLs also exist^8–10^, making ligand specificity exceptionally challenging to predict from sequence alone.

Existing tools are poorly suited to closing this gap. Traditional docking programs (e.g. Vina^11^ or GALigandDock^12^) struggle because receptor flexibility and the high hydrophobicity of AHL ligands confound physical scoring functions. Purely sequence-based or machine-learning approaches are limited by data scarcity: only about 130 LuxR-LuxI systems have an experimentally defined preferred AHL^13,14^ and only 19 co-crystalized structures of unique AHL-LuxR complexes exist in the RCSB Protein Data Bank (PDB), compared to the wide diversity (>10,000) of LuxR homologs in genomic databases (Wellington Miranda et al.^13^ and this work).

Deep learning co-folding prediction models offer a promising alternative, since they are trained to predict the structure of any protein-ligand complex given the protein sequence and ligand specification^15–17^. The constrained AHL ligand space (about 60 described or predicted molecules^14^; Supplemental Table S1) further makes LuxR-AHL specificity a tractable testbed for evaluating these models.

We ensembled multiple protein-ligand co-folding models into a single pipeline which we name COPAL (combining ordered predictions of audited ligands) to robustly model LuxR-AHL interactions (Figure 1d). COPAL outperforms existing methods at ranking LuxR-AHL associations on a leakage-controlled benchmark, and we show its practical value by nominating candidate AHLs for three previously uncharacterized LuxR homologs.

To broaden the QS research community’s access to COPAL predictions, we release here the ranked AHL-LuxR prediction hub (RALPH), which provides COPAL rankings for 10,496 unique LuxR homologs against our AHL ligand pool. We pair RALPH with RALPH-BLAST, a lightweight method that extends rankings to LuxR homologs outside the hub and recovers most of COPAL’s performance on the RefAHL benchmark^14^. Because COPAL combines model rankings without any task-specific training, the same framework extends to other receptor-ligand systems where a defined candidate pool is available.

## Results

### A structure-based ensemble of protein-ligand co-folding models enriches preferred LuxR-AHL interactions in top-ranked predictions

COPAL combines four independent rankings drawn from three protein-ligand co-folding models: AlphaFold3 (AF3)^15^, RosettaFold3 (RF3)^17^, and Boltz2^16^. For each model we rank ligands by interface predicted template modeling (ipTM) scores, adding a fourth ranking from Boltz2’s affinity prediction. We then merge all four into a single consensus ranking, weighting each equally (Figure 1d).

We benchmarked COPAL using a set of 96 LuxR-homologs with their experimentally defined AHL ligand (referred to as “preferred AHL”, or AHL_pref_, in this work) compiled from our recently published RefAHL database^14^ (Supplemental Table S2). This set is “leakage-free”, that is: LuxRs whose preferred AHL pairings could not be trivially inferred from a closely related, ligand-bound structure already present in the co-folding models’ training data (selection criteria in METHODS)

For each receptor in this set, we ran the COPAL pipeline against an audited pool of 58 candidate ligands (57 AHL compounds plus one non-AHL decoy HEHEAA; pool composition in METHODS) generating protein-ligand co-folding predictions with three independent models and the Boltz2 affinity predictor, then integrated their confidence metrics into a single ensemble ranking (Figure 1d; full per-receptor rankings in Supplemental Table S3). We evaluated performance against the experimentally defined preferred AHL (AHL_pref_) using mainly top-K enrichment, defined as the fraction of receptors for which AHL_pref_ was ranked among the K highest-scoring candidates.

Across this benchmark, COPAL consistently enriched AHL_pref_ among top-ranked candidates: the four-model ensemble achieved the highest top-1, top-3 (best 5% of pool), top-6 (best 10%), and top-10 (best 17%) enrichment rates of any method tested, alongside the lowest median and mean ranks and the highest mean reciprocal rank (Table 1, Figure 2a). These results indicate that integrating complementary model representations improves inference of LuxR ligand specificity beyond any single predictor. No individual co-folding model matched COPAL on any enrichment metric, indicating that the models make partially independent errors that ensemble integration corrects. As a baseline, we also implemented a method that predicts a receptor’s ligand ranking purely from sequence similarity to receptors of known specificity, which we refer to as Seq-ID (see METHODS). COPAL outperforms Seq-ID on all enrichment metrics except for a small 1% difference in top-1 success rate (Table 1 and Figure 2a).

**Figure 2.**
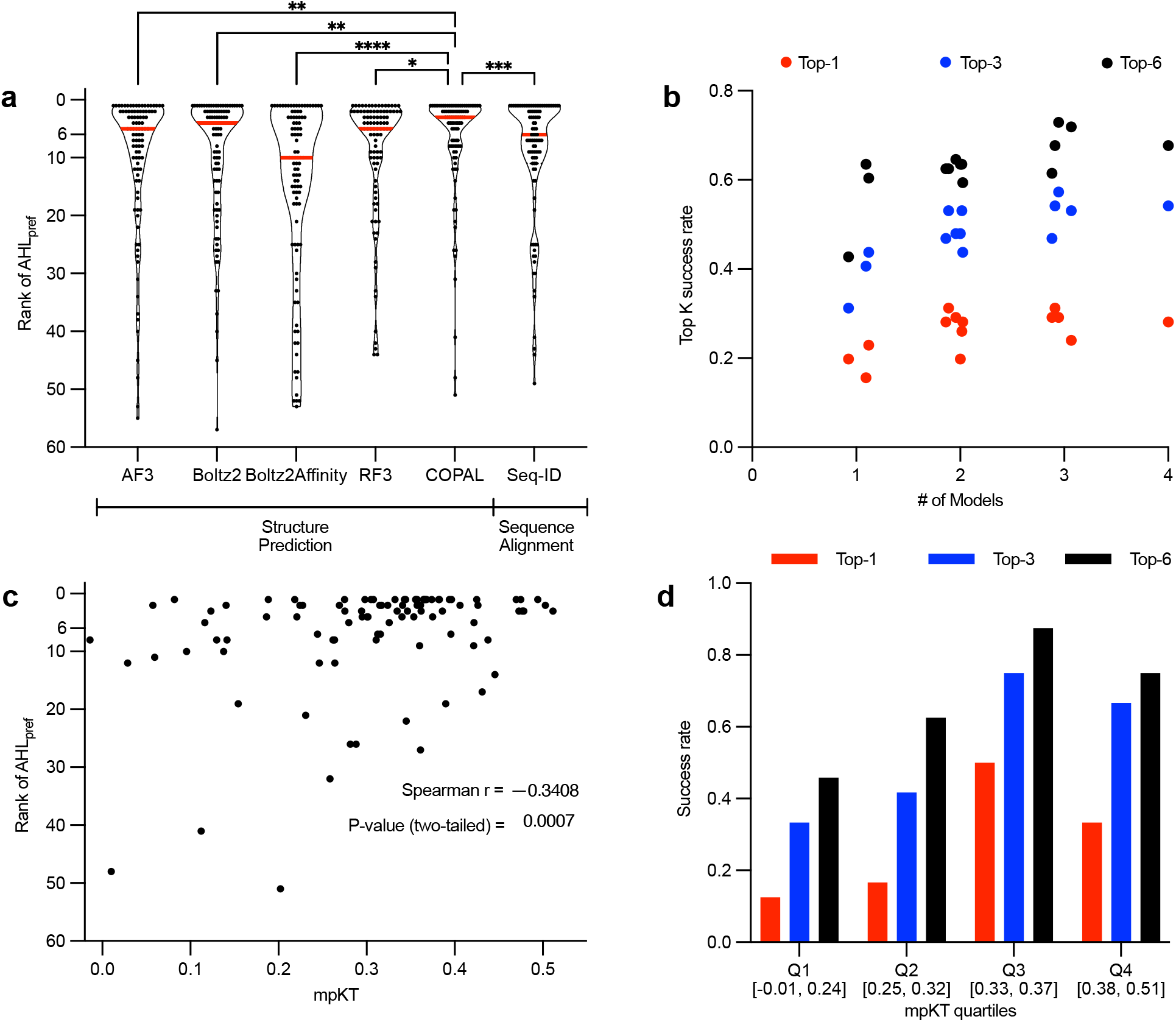
COPAL outperforms individual co-folding models for improved enrichment of AHL_pref_ on LuxR rankings and provides an interpretable confidence metric. **a)** The rank assigned to the cognate AHL ligand (AHL_pref_) by each model when tested against a pool of 58 possible AHLs (n=96 LuxR homologs per group). The y-axis is inverted so that lower rank values (better performance) appear at the top. The red lines indicate medians. Statistical comparisons between COPAL and each individual model were performed using a Friedman test with Dunn’s multiple comparisons post-hoc test (matched by target), with COPAL set as the control column. Adjusted p-values are indicated as: ns p≥ 0.0332, * p < 0.0332, ** p < 0.0021, *** p < 0.0002, **** p < 0.0001. **b)** Top-6, Top-3, and Top-1 success rates for all possible arrangements of individual models included in ensembles of different sizes. **c)** Scatterplot of COPAL’s ranking for the preferred AHL versus mpKT (mean pairwise Kendall’s tau concordance across individual models), a metric of inter-model agreement on the ligand ranking. **d)** Top-1, Top-3, and Top-6 success rates binned by mpKT quartile ranges.

**Table 1.** Summary metrics of the performance of the individual models and COPAL on the RefAHL benchmark. . Each row reports the ligand-ranking performance of one method evaluated on the leakage-free RefAHL benchmark (n = 96 LuxR-AHL pairs): the three protein-ligand co-folding models (AF3, Boltz2, RF3), the Boltz2 affinity head (Boltz2Affinity), the sequence-identity nearest-neighbor baseline (Seq-ID), and the full four-model COPAL ensemble. Columns give the median and mean rank assigned to the preferred AHL (AHL_pref_) among the 58 candidate ligands; the Top-1, Top-3, Top-6, and Top-10 success rates (the fraction of receptors for which AHL_pref_ fell at or above the indicated rank); and the mean reciprocal rank. Arrows in the header indicate the direction of better performance (↑ higher is better; ↓ lower is better). Bold and red marks the best value and underline the second-best value within each column. COPAL achieves the best, or tied-best, score on every metric except Top-1 success rate, where it trails the Seq-ID baseline by about 1%.

| Model | Median rank (↓) | Mean rank (↓) | Top-1 success rate (↑) | Top-3 success rate (↑) | Top-6 success rate (↑) | Top-10 success rate (↑) | Mean reciprocal rank (↑) |
| --- | --- | --- | --- | --- | --- | --- | --- |
| AF3 | 5 | 10.36 | 18.8% | 39.6% | 56.3% | 70.8% | 0.345 |
| Boltz2 | <u>4</u> | 9.74 | 22.9% | <u>43.8%</u> | 60.4% | 67.7% | 0.378 |
| Boltz2Affinity | 10 | 15.13 | 19.8% | 31.3% | 42.7% | 52.1% | 0.299 |
| RF3 | 5 | <u>9.26</u> | 15.6% | 40.6% | <u>63.5%</u> | <u>74.0%</u> | 0.34 |
| Seq-ID | 6 | 9.5 | **29.2%** | 42.7% | 52.1% | <u>74.0%</u> | <u>0.407</u> |
| **COPAL** | **3** | **6.99** | <u>28.1%</u> | **54.2%** | **66.7%** | **82.3%** | **0.45** |

Ablation analysis across all model subsets (Figure 2b; Supplemental Table S4) confirmed that multi-model ensembles consistently outperform individual models, and that adding more models generally improved enrichment. A three-model subset (Boltz2 + Boltz2Affinity + RF3) showed the highest top-6 enrichment overall: top-6 success rate of 73% vs 68% for the four-model ensemble. However, we adopted the full four-model ensemble for COPAL because the difference was small, the validation set is limited (n=96), and the additional model (AF3) has been shown to generalize better to protein-ligand interactions under-represented in training data^18^, which is the regime most relevant to the uncharacterized LuxR homologs COPAL is intended to characterize.

### Agreement between models provides an interpretable measure of prediction reliability of LuxR-AHL interactions

COPAL enrichment varied substantially across receptors, motivating a per-receptor ranking confidence value. We used ensemble-wise inter-model agreement, quantified as mean pairwise Kendall-Tau rank correlation (mpKT) across the rankings output by all pairs in the four-model ensemble, as a metric of how confident COPAL is in each per-receptor ranking (Figure 2c). Higher mpKT correlated with better placement of AHL_pref_ [Spearman ρ =-0.3408 (95% CI:-0.5111 to-0.1447), p (two-tailed) = 0.0007, n = 96)], and receptors in the top half of observed mpKT (mpKT >0.3257) were predicted correctly at substantially higher rates than those in the bottom half (Figure 2d). Inter-model agreement therefore provides interpretable confidence for prioritizing COPAL rankings for experimental follow-up. In contrast, mean individual model ipTM or affinity probability scores over the top-6 ranked ligands are not significantly correlated or substantially less correlated with AHL_pref_ enrichment on top positions in their rankings (Supplemental Figure S1).

### COPAL performance depends on preferred AHL molecular features

While the exact AHL_pref_ was not always enriched in the top-6 positions, the ensemble frequently prioritized chemically similar ligands. To quantify this effect, we computed the structural similarity (MCS-based Jaccard similarity; see METHODS) between the top-k ranked set of ligands and the experimentally determined AHL_pref_ (Figure 3a). Among top-1 predictions, the median structural similarity to AHL_pref_ was 0.8536, and 60% of top-1 predictions had structural similarity ≥ 0.8. This increased to 88% and 93% when considering the highest similarity on the top-3 or top-6 ranked candidates, respectively (n=96). These results indicate close structural similarity between predicted and known ligands even when the exact match was not top ranked. More broadly, structurally similar AHLs may also bind a given LuxR homolog, and there is generally no reason to assume otherwise^19^.

**Figure 3.**
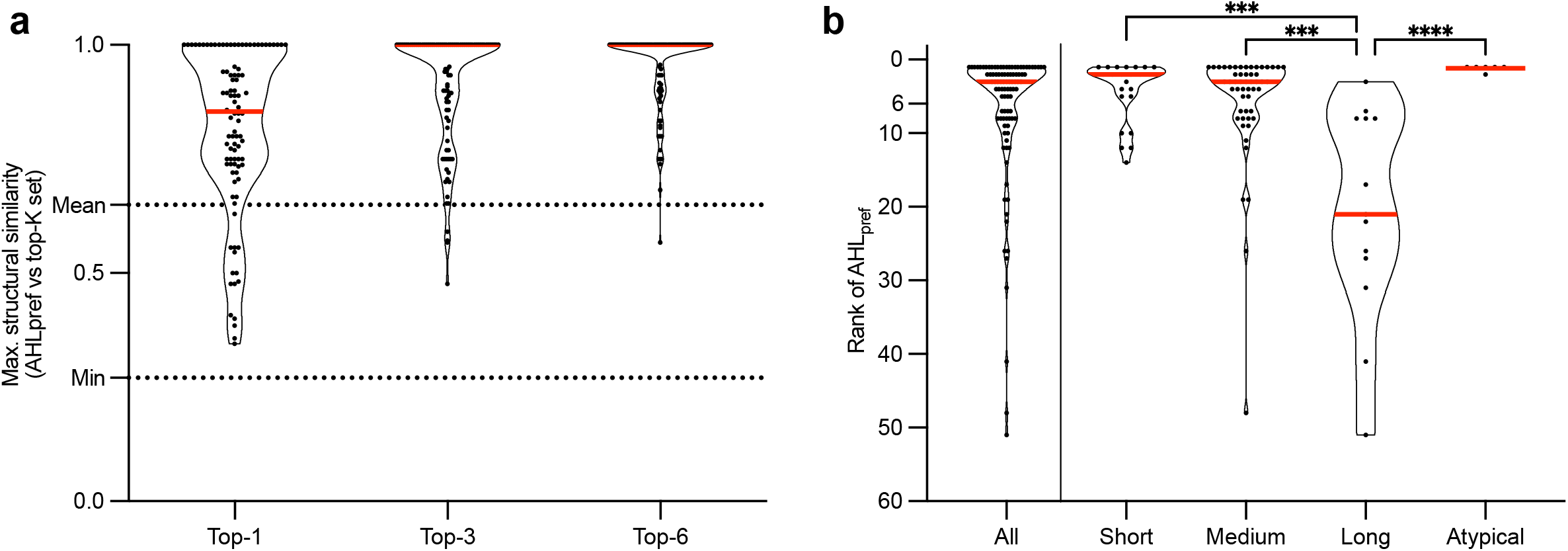
COPAL enriches ligands similar to AHL_pref_ at top positions, but under-performs when AHL_pref_ has a long chain acyl tail (14 or more carbons). **a)** Maximum Common Substructure (MCS) Jaccard similarity between AHL_pref_ and the highest-ranked AHL within the Top-1, Top-3, and Top-6 ranked positions. Each circle represents the maximum MCS similarity for a given LuxR homolog. The dashed horizontal lines indicate the mean and the minimum pairwise MCS similarity across all possible AHL-AHL pairs in the pool. **b)** Distribution of COPAL ranks for AHL_pref_, stratified by acyl-chain category: Short (4–6 carbons), Medium (8–12 carbons), Long (14 or more carbons), and Atypical (structurally non-standard acyl chains, e.g., aromatic; see Supplemental Table S1). The combined distribution for every acyl-chain category is shown to the left of the vertical line on category (labeled “All”). Kruskal-Wallis test (H=24.11, p<0.0001, n=103 across the Short, Medium, Long, and Atypical groups), followed by Dunn’s multiple comparisons post-hoc with Long set as the control category was performed. COPAL ranks for the Long category (n=14) were significantly worse than every other category: Long vs. Short (n=29; adjusted p=0.0008, ***), Long vs. Medium (n = 54; adjusted p = 0.0008, ***), and Long vs. Atypical (n = 6; adjusted p < 0.0001, ****). Significance thresholds as in Figure 2.

Separating rankings by the acyl-chain length of the AHL_pref_ (Figure 3b) revealed that COPAL performed consistently for short (4-6 carbons; n = 25, median rank = 2), medium (8-12 carbons; n = 52, median rank = 3), and atypical AHLs (aromatic and branched chain AHLs; n = 6, median rank = 1), but degraded sharply for long-chain AHLs (≥14 carbons; n = 13, median rank = 21). Among individual models, Boltz2 exhibited a strong bias toward smaller ligands whereas AF3 and RF3 showed more moderate size dependence (Supplemental Figure S2a to S2d). In contrast, Boltz2Affinity predictions displayed minimal correlation with ligand size and retained consistent performance for larger AHLs (Supplemental Figure S2d). These observations define the regime in which COPAL predictions are most reliable and highlight limitations imposed by the distribution of protein-ligand complexes represented in current structural datasets used to train protein-ligand co-folding models.

### COPAL resolves subtle differences in LuxR ligand specificity

As a few amino-acid changes in specific selectivity residues in a LuxR-homolog can produce pronounced shifts in the preferred LuxR ligand^13^, we tested COPAL’s sensitivity by comparing predictions for wild-type *Pseudomonas aeruginosa* LasR and a previously characterized three-substitution variant, LasR^L125F,A127M,L30F^. For wild-type LasR, COPAL rankings are enriched for 12-carbon acyl chain ligands, and its preferred AHL, 3OC12-HSL, is ranked in the top six (Supplemental Table S3 and Figure 4a). Relative to wild-type LasR, the rationally designed variant LasR^L125F,A127M,L30F^ shows increased responses to 3OC10-HSL and broader responsiveness to shorter-chain AHLs (Wellington Miranda et al^13^ and Figure 4b) – changes that are captured by the COPAL rankings (Figure 4a). By contrast, the baseline sequence-identity predictor assigns both LasR and LasR^L125F,A127M,L30F^ to the same closest sequence in the leaked-RefAHL database, and therefore fails to capture the subtle differences in ligand specificity. We note that COPAL did not capture all of the LasR^L125F,A127M,L30F^-AHL ligand activities, as 3OC12-HSL remains equally active as 3OC10-HSL^13^ (Figure 4b), even though its COPAL ranking is reduced (Figure 4a). However, these results suggest that COPAL structure-based ranking can resolve LuxR ligand specificity at a resolution relevant to biological signaling.

**Figure 4.**
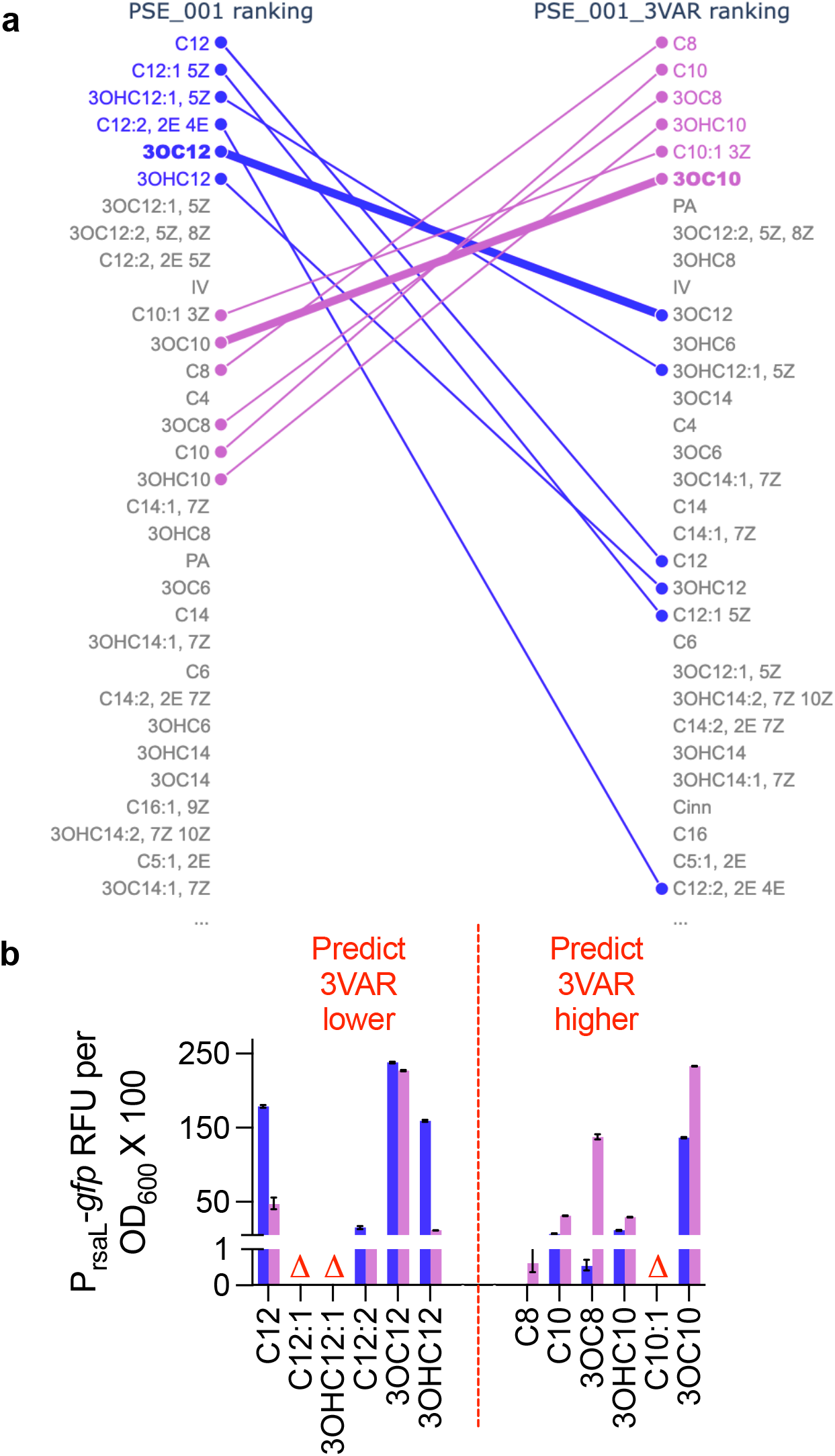
COPAL rankings capture ligand specificity changes in a rationally engineered LuxR homolog variant. **a)** Bump plot comparing the top COPAL rankings for LasR (Pse_001, blue color) vs. LasR^L125F,A127M,L30F^ (Pse_001_3var, lilac color), which harbors 3 amino acid changes to match analogous residues in MupR (Pse_008) as described in the text. Bolding indicates the AHL_pref_ for LasR (3OC12) and MupR (3OC10). COPAL rankings are ordered with the best (top-1) at the top of the list. **b)** Activity of LasR (blue) and LasR^L125F,A127M,L30F^ (purple) when expressed in *E. coli* grown in the presence of the indicated AHL compounds (1 μM, x-axis). COPAL predicts the AHL compounds to the left of the red dashed line would have lower 3VAR activity compared to LasR, and compounds to the right would have higher activity. Red triangles (D) indicate the AHL compound is not commercially available.

### COPAL identifies previously uncharacterized LuxR-AHL interactions

Encouraged by the COPAL results above, we set out to evaluate its performance on uncharacterized LuxR-AHL interactions. We first examined *Mesorhizobium* sp. NJ3^20^, which has a QS LuxR-homolog system with high sequence similarity (81% identity) to one recently described in *Mesorhzobium japonicum* MAFF 303099. The AHL_pref_ for MAFF 303099 is an unusual signal, 2E,4E-dodecadienoyl-HSL (C12:2-HSL), and requires an accessory crotonase enzyme for its synthesis^21^. Generally, I-R homologs sharing this high level of identity use the same AHL signal^22^, but because the NJ3 strain lacks a crotonase gene, we hypothesized that NJ3 used a different AHL ligand than MAFF 303099. Our COPAL results supported this prediction, as the NJ3 LuxR top-ranking AHL was C8-HSL and the MAFF-produced AHL, C12:2-HSL, ranked 15^th^ (Supplemental Table S3). Consistent with COPAL rankings, we found the major AHL present in NJ3 culture extracts was indeed C8-HSL (Supplemental Figure S3a). As confirmation of these results, we measured transcripts of a non-coding RNA reported as QS-regulated in a related strain^23^ and confirmed C8-HSL activated its expression in a LuxR-dependent manner (Figure 5a). We next turned to an undefined LuxR homolog (named TuxR here) in the phototrophic *Rhodopseudomonas palustris* TIE-1^24^. To assess our COPAL predictions, we created a *R. palustris* TuxR-dependent reporter (Figure 5b) and surveyed the AHL inventory (Supplemental Figure S3b). We found C10:1-HSL (top-6 by COPAL) was the major AHL produced by the cognate TuxI, with lesser amounts of C10-and C8-HSL (COPAL ranked 2 and 5, respectively) also detected. All of the top-5 ranked AHLs had TuxR activity (Figure 5b), with C10-HSL (COPAL top-2 ranked and a minor AHL product of the cognate TuxI synthase) showing the most activity. Unfortunately, the major AHL, C10:1-HSL (top-6 ranked), is not commercially available for testing.

**Figure 5.**
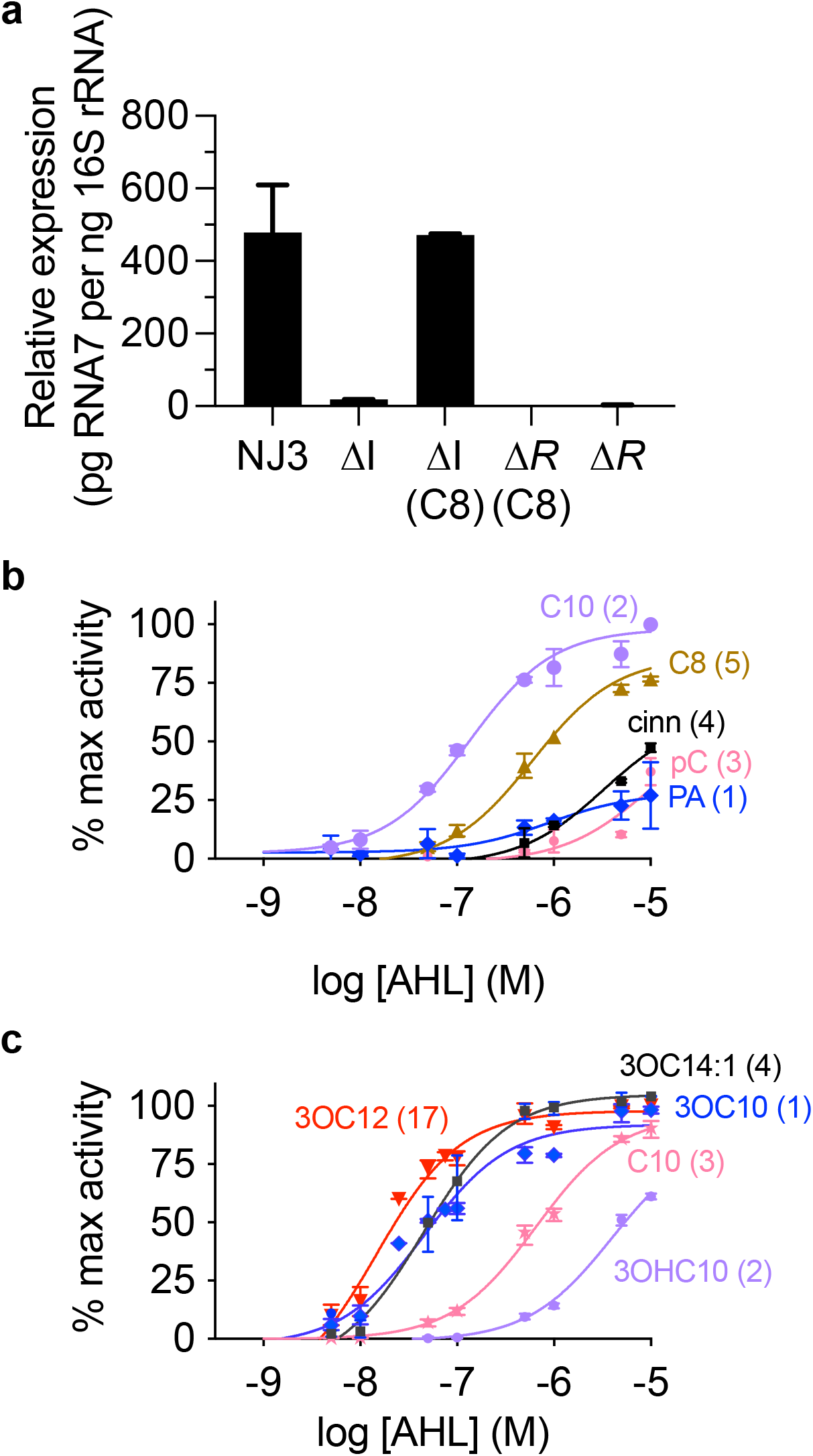
COPAL predicts active AHL ligands for previously uncharacterized LuxR homologs. **a)** Expression of non-coding RNA element [RNA7, ^21^] is dependent on its LuxR homolog and the presence of C8-HSL in *Mesorhizobium* NJ3 strains. We added 50 nM of C8-HSL to either the Δ*luxI1 (*Δ*I*) or Δ*luxR1 (*Δ*R*) mutant where indicated. **b)** Activity of *R. palustris* TuxR expressed in the heterologous host *P. putida* with increasing doses of select AHLs. **c)** Activity of *S. odorifera* LuxR homolog (SO_1110) expressed in the heterologous host *P. putida* with increasing doses of select AHLs. In **b** and **c**, the predicted COPAL ranking for each AHL are indicated in the parentheses.

Finally, we examined a third undefined LuxR homolog in *Serratia odorifera* DSM 4582^25^ (SO_1110), a strain isolated from human sputum. Metabolomics analysis of *S. odorifera* extracts demonstrated that its cognate LuxI produced 3OC12-HSL as its major product, with lesser amounts of 3OC10-and 3OC14-HSL (Supplemental Figure S3c). We were surprised to find that 3OC12-HSL ranked poorly by COPAL – only 17^th^. We created a reporter for this LuxR homolog and we found that the 3OC12-HSL dose-response curve was nearly indistinguishable from those of 3OC10-HSL (COPAL rank 1) and 3OC14:1 (COPAL rank 4) as seen in Figure 5c. Taken as a whole, our experiments using three LuxRs from diverse bacterial strains support the conclusion that, even with some limitations, the COPAL pipeline provides useful information for predicting LuxR-AHL ligand interactions.

### A community resource for LuxR functional annotation

Although COPAL provides an accurate method for modeling LuxR-AHL interactions, its computational requirements (several GPU hours for a single ranking) limit routine use by many laboratories. To enable broad access to these predictions, we applied the COPAL pipeline to a large, non-redundant collection of about 10k LuxR homologs curated from the JGI IMG metagenomics database (see METHODS) and compiled the results into the ranked AHL-LuxR prediction hub (RALPH).

While RALPH covers most of the LuxR sequence space, newly sequenced LuxR receptors or variants with substantial sequence divergence may not be represented. To address this limitation, we developed a simple pipeline on top of the RALPH resource that converts it to a BLAST database that can be quickly queried against any LuxR protein sequence. Rankings of BLAST hits at high identity cutoffs are then averaged and used as the predicted AHL ranking for the new LuxR sequence. When compared to COPAL, RALPH-BLAST enriched experimentally associated ligands among top-ranked predictions with a similar performance (Figure 6), validating RALPH-BLAST as a resource for studying QS systems in metagenomics databases and newly sequenced LuxRs.

**Figure 6.**
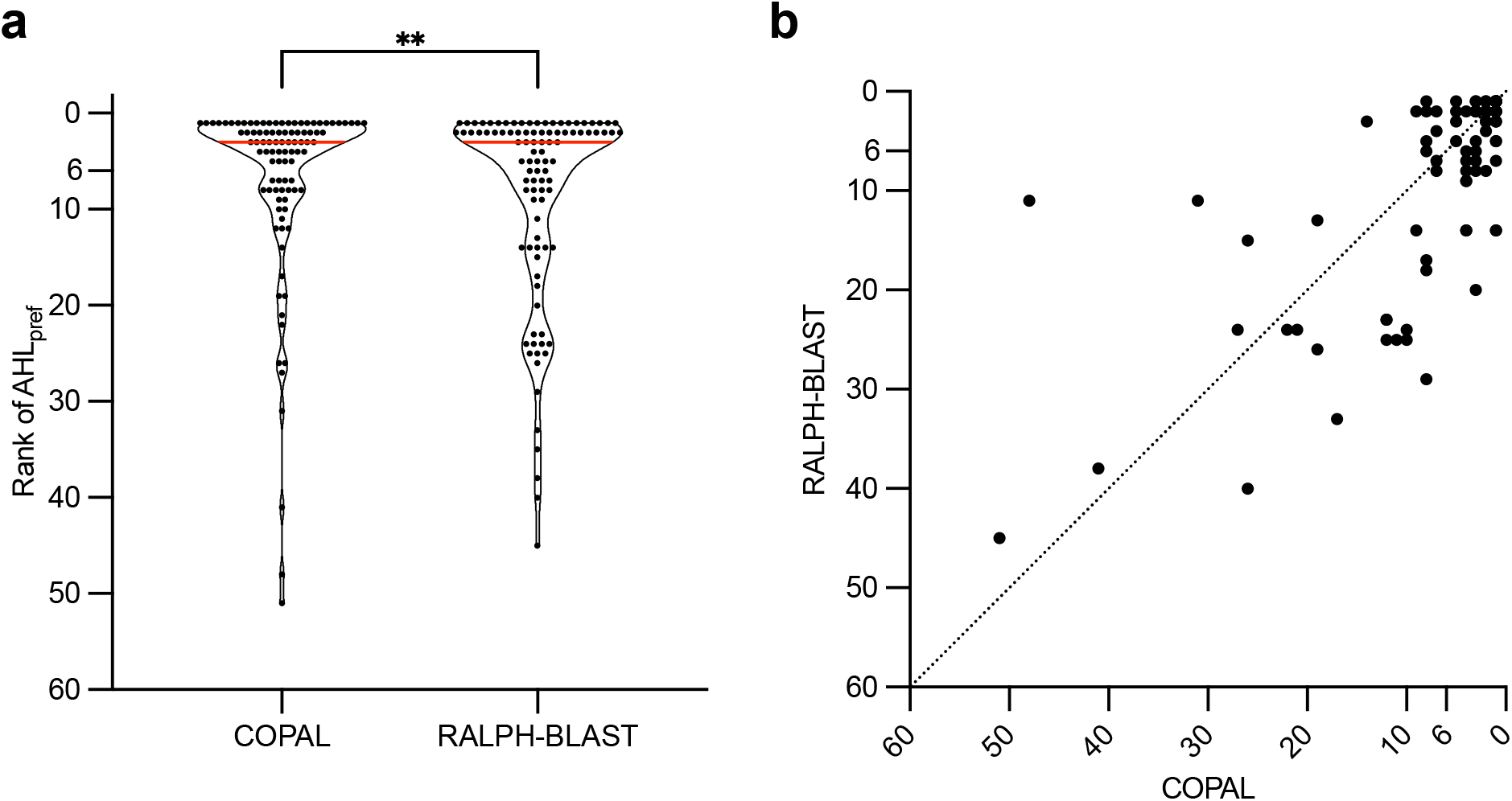
A simple predictor built on top of RALPH can distill COPAL’s performance on the RefAHL benchmark set. **a)** Violin plot comparing the rank distribution of AHL_pref_ between the full COPAL pipeline and the RALPH-BLAST approach across the benchmark set. Red line in each violin indicates the median. RALPH-BLAST achieves a rank distribution similar to COPAL (Spearman r=0.71 between paired AHL_pref_ ranks, p<0.0001), recapitulating most of COPAL’s performance. COPAL retains a small but significant advantage in paired comparison (Wilcoxon matched-pairs signed-rank test, p=0.0058, n=96, 25 tied pairs). **b)** Scatterplot of RALPH-BLAST predicted rank versus COPAL predicted rank of AHL_pref_ for each LuxR homolog. The dashed line indicates x = y (perfect agreement). Most points cluster near the origin, indicating concordance between the two methods, with RALPH-BLAST occasionally assigning higher (worse) ranks to cases where COPAL also struggles. Significance thresholds as in Figure 2.

### COPAL can be extended beyond AHL chemistry

Some LuxR homologs present in sequence databases, and therefore in RALPH, are not AHL-responsive. The PipR/OryR subfamily is one such case. These receptors have evolved to respond to structurally distinct, non-AHL ligands present in plant host exudates^10,26^. To test whether COPAL extends beyond canonical AHL chemistry, we added HEHEAA as a 58th candidate ligand (rationale and methodology in METHODS). COPAL ranked HEHEAA #1 for PipR homologs encoded by *Pseudomonas* sp. GM79 (PipR_GM79) and *Pseudomonas syringae* DC3000 (PipR_DC3000), while all leakage-free RefAHL benchmark LuxRs ranked HEHEAA at or above position 34, indicating that the structure-based ranking captures general features of LuxR ligand recognition and can extend to non-AHL signals. Because HEHEAA is itself an ethanolamine derivative, a top HEHEAA ranking should serve as a positive indicator that the receptor binds ethanolamine-derived ligands more broadly^26^, as additional ethanolamine-derived related signals are thought to exist^26,27^. HEHEAA was added as part of the ligand pool during RALPH construction.

## Discussion

COPAL shows that an unweighted ensemble of protein-ligand co-folding models predicts LuxR ligand specificity more reliably than any single model, placing the preferred AHL among the top-6 of 58 ligand candidates for 68% of receptors on a leakage-free benchmark. COPAL resolved the specificity shift of an engineered LasR variant and correctly nominated C8-HSL for the previously uncharacterized *Mesorhizobium sp.* NJ3 receptor, which we confirmed experimentally. Packaged as the RALPH resource and its lightweight BLAST surrogate, these predictions are now available across roughly 10,000 LuxR homologs, providing the quorum sensing community a practical tool to prioritize experiments and, more broadly, a template for applying co-folding ensembles to other receptor-ligand families with a defined candidate pool. Naturally, experimental validation of COPAL predictions for previously uncharacterized LuxR receptors, prioritized via the RALPH resource, is the immediate next step for future work.

However, several limitations should be noted. First, COPAL performance exhibits a systematic dependence on AHL acyl-chain length, with significantly reduced enrichment for longer-chain ligands. This effect may reflect biases in the structural training data available to current co-folding models, which contain fewer examples of protein-ligand interactions with extended acyl chains. Second, the candidate ligand pool is restricted to 57 AHL and 1 non-AHL molecules; LuxR receptors that recognize structurally unrelated signals (other than HEHEAA) or not-yet-characterized AHLs would not be captured by the current pool. Third, while inter-model agreement (mpKT) provides a useful reliability indicator, substantial variance in ranking accuracy persists even at moderate agreement levels (Figures 2c and 2d), indicating that mpKT alone is not sufficient to guarantee prediction quality for any individual receptor.

A central methodological contribution of this work is that aggregating the ranked outputs of several co-folding models, without any learned weighting, yields a more reliable predictor than any single model. We chose unweighted rank summation over a learned weight ensemble because, with fewer than 100 leakage-free LuxR-AHL pairs available for fitting, data-driven weighting would risk overfitting and generalize poorly. Unweighted aggregation also requires no additional training and applies to any new co-folding model that outputs protein-ligand confidence scores. This provides a general formula for prioritizing candidate ligands in other receptor-ligand systems where labeled data are scarce, but a defined candidate pool is available, and the framework should gain accuracy as the underlying co-folding models improve. Rather than replacing experimental validation, COPAL is intended to accelerate discovery by focusing experimental effort on the most plausible signaling interactions, ultimately contributing to a more complete understanding of bacterial communication networks and their roles in microbial ecology, evolution, and host interactions.

## Methods

### Audited ligand pool

Candidate ligands consisted of an audited pool of 58 compounds: 57 AHL molecules experimentally observed^14^, or predicted to exist^28^, in LuxI-LuxR type quorum sensing systems; plus N-(2-hydroxyethyl)-2-(2-hydroxyethylamino) acetamide (HEHEAA) as the sole non-AHL member^10,26^. HEHEAA essentially functions as a non-AHL decoy in RefAHL benchmarking and as a prioritization target for potential non-AHL-responsive receptor homologs (PipR-family) with no modification to the pipeline architecture. Ligands were represented using isomeric molecular SMILES strings to preserve stereochemistry and we indicate the compounds that are commercially available (Supplemental Table S1). Note that there are likely not-yet-discovered AHL signal structures present in nature which, by definition, are not included in our COPAL pool of audited ligands. For example, we did not include all possible isomerization states or locations for AHLs with double bonds.

### Sequence identity analysis and leakage-free benchmark curation

Structures of quorum-sensing LuxR-family transcriptional regulators in complex with ligands were retrieved from the RCSB Protein Data Bank (PDB) using the RCSB Search API. The “LuxR family” was defined as the subset of LuxR-type proteins bearing the N-terminal autoinducer-binding domain characteristic of acyl-homoserine lactone (AHL) receptors, corresponding to Pfam family PF03472 (“Autoind_bind”) and the equivalent InterPro entry IPR005143 (“Transcription factor LuxR-like, autoinducer-binding domain”). We filtered the returned entries by release date earlier than January 1^st^ 2024, which is the inclusion criteria for the training set of RF3 (RF3 had the latest training set cutoff date). In total, 51 unique PDB entries fit these criteria. Only 22 PDB crystal structure entries of LuxR-AHL interactions were represented in the 51 PDB entries (Supplemental Table S5), the rest being non-canonical AHL ligands, crystallization aids, NMR structures, or non-AHL ligands. These were spread over nine unique LuxR sequences, which were identified as potential training-set leakage sources (Supplemental Table S5). When multiple PDB entries contained the same LuxR homolog bound to the same AHL, we retained only those entries representing the canonical receptor-ligand complex and excluded additional entries containing the same LuxR-AHL pair in complex with non-LuxR homolog proteins. For example, 3IX3 and 2UV0 both contain Pse_001 bound to 3OC12-HSL and were retained (Supplemental Table S5); 4NG2, 6V7W, and 6V7X also contain Pse_001with 3OC12-HSL, but in complex with additional proteins, and were therefore excluded.

Seven of these nine LuxR-homologs are themselves RefAHL entries; QscR and SdiA are orphan LuxR homologs included only as reference sequences for the identity analysis. Each of the roughly 130 RefAHL receptors was globally aligned against each of the reference sequences using global Needleman-Wunsch alignment with the BLOSUM62 substitution matrix, a gap opening penalty of −10, and a gap extension penalty of −0.5, as implemented in Biopython (v1.84). Sequence identity was defined as the number of identical residue pairs in the alignment divided by the total alignment length including gaps. Following the 40% sequence identity threshold established by recent structure prediction benchmarks^29^, we classified each non-PDB receptor by its maximum sequence identity to any PDB-containing LuxR. Of the 129 non-PDB RefAHL receptors, 101 shared less than 40% sequence identity with all PDB-containing LuxR receptors. The remaining 28 exceeded the 40% threshold and were further evaluated based on whether they share the same preferred AHL as their closest PDB homolog: 26 receptors shared both high sequence identity (42 to 64%) and the same preferred ligand, and were excluded as potentially confounded by training set memorization, while two receptors exceeded 40% identity but were assigned a different preferred AHL than their closest PDB homolog and were therefore retained. We also omitted cases where multiple LuxR sequences are localized next to the same LuxI and no experimental characterization exists to disambiguate which LuxR responds to the AHL made by the LuxI (six such cases). The resulting leakage-free evaluation subset comprised 96 receptors (94 below the identity threshold plus two above threshold with different ligands). BLAST metrics (NCBI BLAST+ v2.11.0, evalue 1e-3, BLOSUM62) were computed in parallel and retained as auxiliary columns in Supplemental Table S3.

### Leveraging an ensemble of protein-ligand co-folding models

Protein-ligand structure predictions were generated using three independent deep learning models: AlphaFold3 (AF3)^15^, RosettaFold3 (RF3)^17^, and Boltz2^16^. These models share a common objective of predicting protein-ligand complex structures from sequence and molecular graph representations but differ in architecture, training regimes, and data curation strategies. Boltz2 affinity predictions were additionally included as an orthogonal signal reflecting predicted binding likelihood rather than structural confidence.

Each model was run independently for every LuxR-ligand pair. For each run, five structural samples were generated using a single random seed, and the prediction with the highest protein-ligand interface confidence (ipTM) was retained for downstream analysis. ipTM was extracted from the model output JSON for AF3 and Boltz2, and from the CSV output for RF3.

For each predicted LuxR-ligand complex, model-specific confidence metrics describing protein-ligand interface quality were extracted. These included interface-level confidence measures derived from predicted pairwise error or distance estimates, as provided by each model. Confidence metrics were used solely for relative ranking of candidate ligands for a given LuxR receptor and were not compared across different LuxR proteins.

### COPAL ranking procedure

COPAL ranks the *N* candidate ligands for each LuxR receptor *p* independently, in three steps. First, multiple predictions per receptor-ligand pair were pooled by their maximum confidence (ipTM) or affinity score (Boltz2 Affinity Probability). Then, within each receptor, every model ranks ligands by descending score (highest is ranked first), with tied scores assigned the most generous (minimum rank):

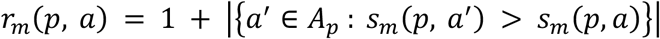

Where *r_m_*(*p*, *a*) is the per-model *m* rank of ligand *a* for receptor *p*, *A_p_* is the panel of *N* candidate ligands, and *s_m_*(*p*, *a*) is the pose-pooled per-model score

Finally, the per-model ranks are aggregated by a parameter-free rank-sum, and the COPAL rank is ascending in this aggregate score (lowest is ranked first):

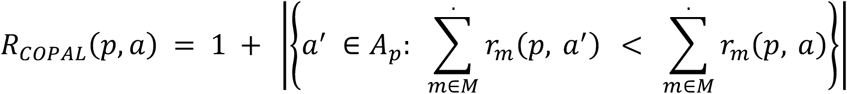

When a strict ordering was required, any remaining ties in the per-model or COPAL ranks were broken randomly.

### Structural similarity metric

To assess structural similarity between top-ranked and experimentally associated ligands, we implemented a similarity metric specific for AHL-family of ligands by using maximum common substructure (MCS)-based Jaccard index over heavy atoms. The MCS was identified with RDKit’s FindMCS using element-aware atom matching, ring atoms constrained to match ring atoms, and bond-order-agnostic bond matching (rdFMCS.BondCompare.CompareAny). To retain sensitivity to bond-order differences that distinguish closely related AHL analogs (e.g. 3-oxo versus 3-hydroxy substituents, or saturated versus mono-or di-unsaturated acyl chains), the effective common-atom count was reduced by one for every bond whose order differed between the two molecules within the matched substructure. The reported similarity is the Jaccard ratio of this effective common atom count to the union of heavy atoms across both molecules; hydrogens were excluded throughout.

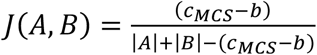

Where *c_MCS_* is the MCS atom count as described above, *b* is the number of MCS bonds whose bond order differs between ligand *A* and *B*, and |*X*| is the number of heavy atoms for any ligand *X*.

### Sequence identity nearest-neighbor baseline

As a reference point which does not depend on structure prediction models, we implemented a baseline that outputs a receptor ligand ranking purely from sequence similarity to receptors of known AHL specificity (which we call Seq-ID). The receptors excluded from the leakage-controlled benchmark (those whose sequences are too similar to a co-crystallized training structure) and that carry an annotated AHL_pref_ were used as a labelled database (which we call Seq-ID database). For each held-out (leak-free) receptor, we computed global pairwise sequence identity (Needleman-Wunsch, BLOSUM62, gap-open-10, gap-extend-0.5) to every Seq-ID database sequence and selected the single most identical receptor as its nearest neighbor. The AHL_pref_ of that neighbor was taken as an anchor ligand, and every other AHL was then scored by its structural similarity to the anchor, ranking ligands in descending order of similarity. Structural similarity was measured using the bond-order-penalized MCS Jaccard similarity-based metric described in Methods. The anchor ligand therefore receives the top rank by construction, with the remaining ligands ordered by chemical proximity to it.

### Inter-model agreement calculation (mpKT)

To assess prediction reliability, agreement between individual model rankings was quantified using Kendall-Tau rank correlation. For each LuxR receptor, pairwise correlations between all model rankings were computed and averaged to yield a mean pairwise Kendall-Tau score (mpKT). This metric was used as an indicator of consensus among models and was analyzed in relation to ensemble ranking performance.

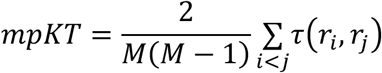

Where *M* is the number of models, *r_i_*and *r*_7_ are rankings for models *i* and *j*, and *τ* is the Kendall-Tau correlation coefficient function.

### Wet-lab experimentation

To assess ligand binding of LuxR homologs, we measured LuxR-dependent QS gene expression. We used the following fluorescent protein-based reporters: *Escherichia coli* pJNL/pJNL^L125F,A127M,L30F^ and pPROBE-P_rsaL_ for Pse_001 (LasR) or Pse_001_3var (LasR^L125F,A127M,L30F^) activity^9,13^; *Pseudomonas putida* pSMC1 for TuxR (encoded by *Rhodopseudomonas palustris* TIE-1) activity (this work, GenBank accession PZ600025) and *P. putida* pSMC2 for the LuxR homolog SO_1110 (encoded by *Serratia odorifera* DSM 4582) activity (this work, GenBank accession PZ60026). Assays were as described previously^13,21^, except that cells were grown in 13-mm boroscillate tubes and transferred to 96-well plates for fluorescence and optical density measurements. For *Mesorhizobium* NJ3_R1 activity, we used reverse transcription quantitative polymerase chain reaction (RT-qPCR) assay to measure transcripts of a non-coding RNA (RNA7) reported to be QS-regulated in a related strain^23^. RNA was prepared as described previously^30^, and 200 ng of total RNA was used to generate cDNA by using the qScript cDNA synthesis kit. Quantitative real-time PCR was performed using 10-and 100-times diluted cDNA product for RNA7 and 16S rRNA, respectively, using the SsoAdvanced Universal SYBR Green Supermix (BIO-RAD). Standard curve for quantification of the NJ3 RNA7 transcripts was generated using dilutions of NJ3 chromosomal DNA and transcript levels were normalized to transcript levels of the NJ3 16S rRNA^31^. For target size and primer sequences see Supplemental Table S7.

To elucidate the AHL signal(s) made by the paired LuxI homologs in *S. odorifera*, *R. palustris* TIE-1, and *Mesorhizobium* sp. NJ3, we grew the indicated strains to stationary phase, extracted cell-free culture fluid twice with acidified ethyl acetate, and analyzed the metabolic profiles using untargeted, high resolution LC-MS/MS (Orbitrap Ascend Tribrid with ZORBAX Eclipse Plus C18 2.1X50, from Thermo Scientific; and a solvent system of water and acetonitrile both containing 0.1% formic acid). Potential AHL products were identified by screening for known and predicted AHLs and potential unknown AHLs in both the MS1 and MS2 modes^28,32,33^. We also analyzed 29 commercially available, chemically synthesized AHL compounds (Supplemental Table S1) to confirm a given AHL’s retention time and MS2 fragmentation pattern.

### Construction of the Ranked AHL-LuxR Prediction Hub (RALPH)

To enable broad access to COPAL predictions, the pipeline was applied to a large, non-redundant collection of LuxR homologs. LuxR homologs were retrieved from the Joint Genome Institute Integrated Microbial Genomes (IMG) database^34^ using the Pfam annotation for the AHL-binding domain (pfam03472). To reduce redundancy, sequences were clustered using MMSeq2^35,36^ with a 90% sequence identity threshold and 0.8 coverage, and sequences shorter than 125 amino acids were excluded. In total, 10,496 LuxR protein sequences were found using this collection strategy and a FASTA file containing their amino acid sequences is provided here (Supplemental File S1 and Supplemental Table S6).

For each LuxR sequence, ensemble rankings and confidence metrics were computed across the candidate ligand pool and compiled into the Ranked AHL-LuxR Prediction Hub (RALPH). RALPH provides precomputed results intended to support hypothesis generation and experimental prioritization without requiring users to run computationally intensive structure predictions. We provide the COPAL results for RALPH in Supplemental Table S6. We also integrated the results into an interactive Google Colab notebook, which allows for homology-based queries of RALPH (https://github.com/davinan/COPAL; DOI: 10.5281/zenodo.21305294).

## Statistical analyses

Statistical analyses were performed using the GraphPad Prism11 software. Unless otherwise specified, all statistics were computed on the leakage-free subset (n = 96) defined in Sequence identity analysis and leakage-free benchmark curation. Top-K success rate is defined as the fraction of receptors for which the preferred AHL ranked at position ≤K among the 58 candidate ligands. COPAL versus individual-model comparisons were tested for significance using Friedman test with Dunn’s multiple-comparisons post-hoc (non-parametric). The association between mean pairwise Kendall-Tau agreement (mpKT) and COPAL preferred-AHL rank was assessed with Spearman’s rank correlation. Differences in AHL_pref_ rank distributions across acyl-chain length categories (Short, Medium, Long, Atypical) were assessed by the Kruskal-Wallis test, a non-parametric one-way analysis of variance appropriate for non-normally distributed rank data. To identify which categories differed from the Long group, a Dunn’s multiple comparisons post-hoc test was performed with the Long category set as the control group, comparing it pairwise to Short, Medium, and Atypical. Adjusted p-values were computed by Prism using Dunn’s correction for the three pre-specified comparisons. Graph Pad Prism thresholds for statistical significance were used: ns p≥0.0332, * p<0.0332, ** p<0.0021, *** p<0.0002, **** p<0.0001.

## Data Availability

The RefAHL benchmark dataset is publicly available via the RefAHL database. Precomputed COPAL rankings comprising the RALPH resource are available at github.com/davinan/COPAL (DOI: 10.5281/zenodo.21305294). Data from the LC-MSMS analyses were deposited as.mzML files at the MassIVE community repository as part of the ProteomeXchange consortium (dataset identifiers MSV000102442, MSV000102444, and MSV000102446) and can be accessed by logging into https://massive.ucsd.edu as MSV000102442_reviewer with the password Reviewer1!. Plasmid DNA sequences were deposited at The National Center for Biotechnology Information (GenBank accessions PZ600025 and PZ600026). All other data supporting the findings of this study are available from the corresponding author upon reasonable request.

## Code Availability

The COPAL pipeline code, RALPH construction, and RALPH-BLAST notebook are available at https://github.com/davinan/COPAL.

## Author Contributions

**Conceptualization**: F.D., A.S., E.P.G., D.N.A. **Methodology**: D.N.A., A.S. **Software**: D.N.A. **Formal analysis**: A.S., D.N.A. **Investigation**: A.S., D.N.A., S.C., B.B., C.M., Y.O. **Data curation:** A.S. **Validation**: A.S., S.C., B.B., C.M., Y.O. **Writing - original draft:** A.S., D.N.A. **Writing - review & editing**: F.D., A.S., E.P.G., D.N.A., S.C., B.B., C.M., Y.O. **Supervision:** F.D., E.P.G. **Funding acquisition:** F.D., E.P.G.

## Funding acknowledgements

F.D and D.N.A are supported by funds from The Defense Threat Reduction Agency (DTRA) (HDTRA1-22-1-0012). This work was supported by NIH grant R35 GM136218 (to E.P.G.). S-M.C. was supported by the National Science and Technology Council Overseas Project for Post Graduate Research (NSTC 114-2917-I-006-004). B.B. was supported by The Fund for Shanxi 1331 Project (1331KSC). We acknowledge the School of Pharmacy’s Mass Spectrometry Center at the University of Washington for providing access to the mass spectrometry instrumentation. We thank Dale Whittington for assistance with protocol development and analysis. Instrument access was made possible in part by NIH grant #S10OD030237. We thank Josh Ramsay for sharing the Bastholm thesis (Ref. 23) in advance of publication.

## Supporting information

Supplemental Talbe 1

## List of Figures, Tables, and Supplementals

Supp File 1. fasta file of ∼10k LuxR protein sequences used in RALPH

Supp Table S1 (.xls) Information/list of compounds in the audited ligand pool (n=58)

Supp Table S2 (.xls) Information/list of the LuxR homologs used in this work (n=138)

Supp Table S3 (.xls) COPAL ranking results table

Supp Table S4 (.xls) Ablation table

Supp Table S5 (.xls) LuxRs with co-crystalized AHL ligands and COPAL rankings table

Supp Table S6 (.xls) COPAL rankings for RALPH

Supp Table S7 (.xls) Primers used for RT-qPCR

## Supplemental Table Descriptions

**Supplemental Table S1. Information and inventory of the audited pool of 58 candidate ligands.** Complete description of the audited ligand pool used as the candidate set for every COPAL prediction: 57 acyl-homoserine lactones (AHLs) observed or predicted in LuxI-LuxR quorum-sensing systems, plus the single non-AHL decoy HEHEAA. Each row describes one compound, with columns for the internal AHL identifier (AHL01–AHL57, and HEHEAA), abbreviation, common name, acyl side-chain category (short, n = 7; medium, n = 16; long, n = 30; atypical, n = 4; non-AHL, n = 1), molecular formula, exact monoisotopic mass, the commercial vendor and catalog number where the compound is available as a standard, and the isomeric SMILES string used to represent stereochemistry in the co-folding models.

**Supplemental Table S2. Information and list of the LuxR homologs used in this work.** Catalog of the LuxR-family receptors compiled for this study, drawn primarily from the RefAHL database together with orphan reference receptors, PipR-family receptors, and the previously uncharacterized homologs now characterized here. Each row describes one receptor, with columns for: category (RefAHL, orphan reference, PipR, or wet-lab characterized), RefAHL confidence rating, RefAHL identifier and published LuxR name, a flag indicating membership in the leakage-free benchmark subset (n = 96), maximum sequence identity to any co-crystallized LuxR–AHL PDB structure, the LuxR amino-acid sequence and length, the experimentally defined preferred AHL (AHL_pref_ id and abbreviation), source bacterial strain or metagenome, PubMed reference IDs (PMID), the paired LuxI homolog name and its IMG/GenBank identifiers, the LuxR IMG/GenBank identifiers, and any relevant notes.

**Supplemental Table S3. Per-receptor COPAL and individual-model ligand rankings on the benchmark set.** Full ranked lists of all 58 candidate ligands for each LuxR receptor in the benchmark set, produced by COPAL and by each individual model. The workbook contains six sheets that share a common layout: COPAL (the four-model ensemble), AF3, Boltz2, Boltz2Affinity, RF3, and Seq-ID (the sequence-identity baseline). For each receptor the leading columns give the LuxR name, category, RefAHL confidence class, the preferred AHL (AHL_pref_ id, abbreviation, and acyl-chain category), the mean pairwise Kendall-Tau inter-model agreement (mpKT), and the rank that method assigned to AHL_pref_; the remaining 58 columns (rank_1…rank_58) list the candidate ligands ordered from best (rank_1) to worst (rank_58). Each receptor occupies a block of three rows that report, in order, the ligand AHL identifier, the ligand abbreviation, and the underlying score at each rank position (the parameter-free rank-sum aggregate on the COPAL sheet; the model’s ipTM or affinity-probability score on the individual-model sheets), followed by a blank separator row. Multiple sheets.

**Supplemental Table S4. Ablation of model combinations in the COPAL ensemble.** Top-K enrichment for every possible subset of the four ranking signals that make up COPAL. Each row corresponds to one of the 15 non-empty combinations of (AF3, Boltz2, Boltz2Affinity, RF3): four Boolean columns indicate which metrics are included and three columns report the resulting Top-1, Top-3, and Top-6 success rates on the leakage-free benchmark (n = 96). The full four-model ensemble adopted for COPAL is the first row, and the three-model subset (Boltz2 + Boltz2Affinity + RF3) gives the highest Top-6 success rate (0.729). **Bold** numbers indicate the best performance and underlined numbers indicate the second best performance over all sets.

**Supplemental Table S5. Co-crystallized LuxR–AHL PDB structures and their COPAL ranks.** List of the RCSB Protein Data Bank entries containing a LuxR-family receptor co-crystallized with an AHL ligand, used to define potential training-set leakage (see METHODS). Each row gives the PDB accession (with the co-crystallized AHL noted in parentheses), a category label (RefAHL, PDB, or orphan), the RefAHL identifier of the LuxR homolog, its experimentally defined preferred AHL (AHL_pref_ id and abbreviation), the AHL actually present in the crystal structure (id and abbreviation), and the COPAL rank assigned to that co-crystallized AHL. Footnotes indicate entries that share the same LuxR homolog bound to a different AHL (grey shading) and entries in which the co-crystallized AHL is not the reported AHL_pref_.

**Supplemental Table S6. Summary of RALPH results: precomputed rankings for 10,496 LuxR homologs.** The Ranked AHL-LuxR Prediction Hub (RALPH): precomputed rankings of the 58-ligand pool for each of the 10,496 non-redundant LuxR homologs curated from the JGI IMG database (sequences provided in Supplemental File S1). Columns give the RALPH identifier (RALPH00000…), the mean pairwise Kendall-Tau inter-model agreement (mpKT), and 58 ranked-ligand columns (rank_1 to rank_58) ordered from best to worst. Each homolog occupies a single row reporting the ligand abbreviation.

**Supplemental Table S7. Primers used for RT-qPCR quantification of the QS-regulated non-coding RNA in *Mesorhizobium* sp. NJ3.** Forward and reverse primer pairs used in the reverse-transcription quantitative PCR (RT-qPCR) assay (Figure 5a) measuring transcript levels of the quorum-sensing-regulated non-coding RNA (RNA7) and of the 16S rRNA normalization control in *Mesorhizobium* sp. NJ3. Columns list the target (NJ3_16S rRNA or NJ3_RNA7), the primer name, the 5′◊3′ oligonucleotide sequence, and the expected amplicon (target) size in base pairs. Nakajima An et al Figure 1

---

**Supplemental Figure S1.**
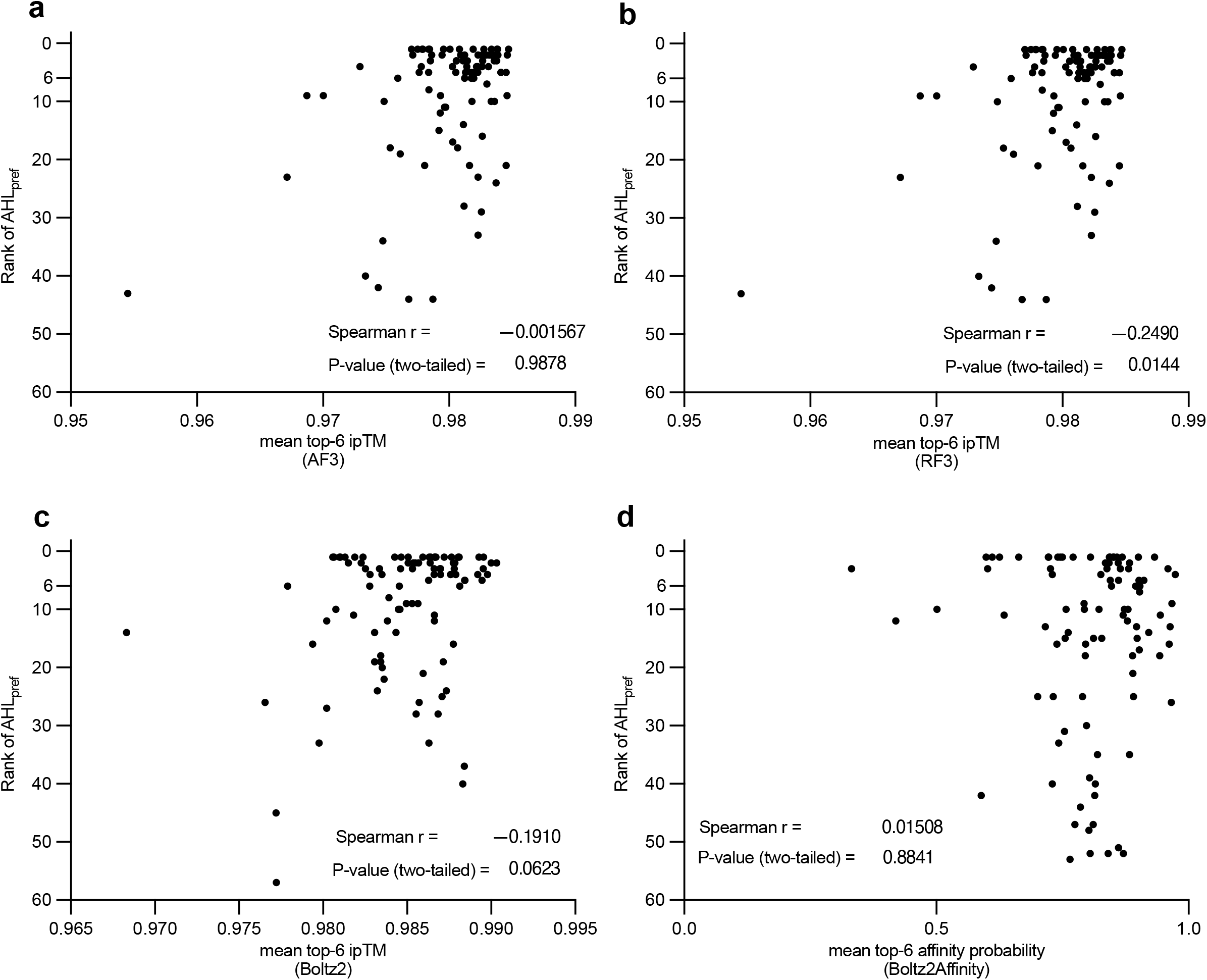
Model-specific aggregated ranking metrics are not predictive of AHLpref enrichment. Ranking performance of individual models plotted as a function of the mean value of the top-6 ranked ligands in the individual model ranking. Spearman r and their p-values annotated in each graph **a)** AF3 ipTM-based ranking. **b)** RF3 ipTM-based ranking. **c)** Boltz2 ipTM-based ranking. **d)** Boltz2 Affinity binary prediction-based ranking. The per-model signals are weak: AF3 r=−0.02 (p=0.99); Boltz2 r=−0.19 (p=0.06); Boltz2Affinity r=+0.02 (p=0.88); RF3 r=−0.25 (p=0.01).

**Supplemental Figure S2.**
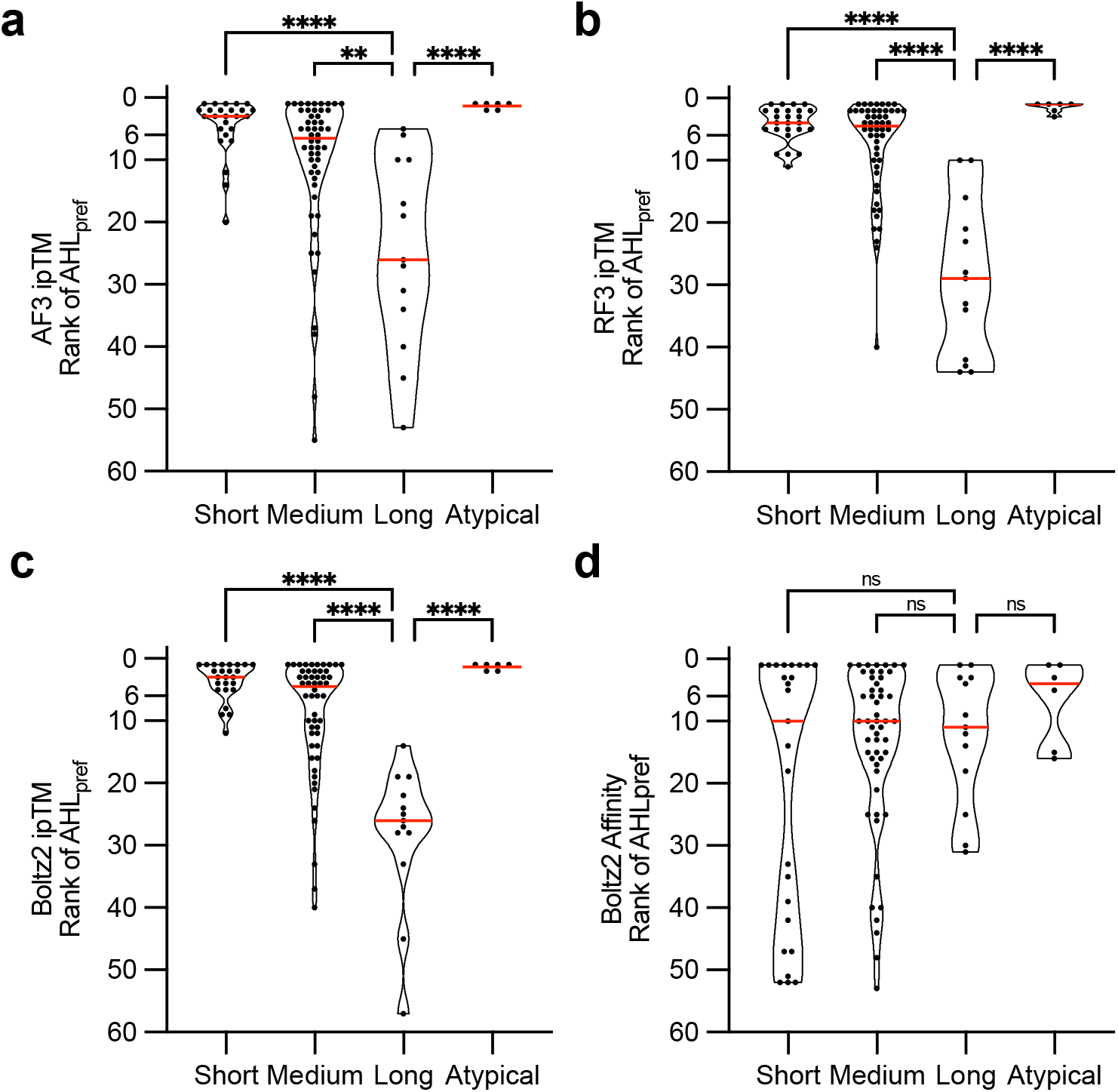
Model-specific sensitivity to AHL molecular size. Ranking performance of individual models within the ensemble plotted as a function of AHL acyl-chain length category (Short, Medium, Long, Atypical) for the AHL_pref_. **a)** AF3 ipTM-based ranking. **b)** RF3 ipTM-based ranking. **c)** Boltz2 ipTM-based ranking. **d)** Boltz2 Affinity binary prediction-based ranking. Red lines in each violin plot indicates the median. AF3 and RF3’s performances show moderate size dependence, Boltz2 ipTM exhibits a much stronger performance bias toward smaller chained AHL_pref_ ligands. Boltz2 Affinity predictions show no statistically significant size dependence (all ns), suggesting more balanced performance across chain lengths. These complementary biases motivate the ensemble approach. Statistical comparisons were performed independently for each panel using Kruskal-Wallis with Dunn’s post-hoc, Long set as the control category. Note that the Atypical group is small (n=6) so statistical comparisons should be interpreted with caution. Significance thresholds as in Figure 2.

**Supplemental Figure S3.**
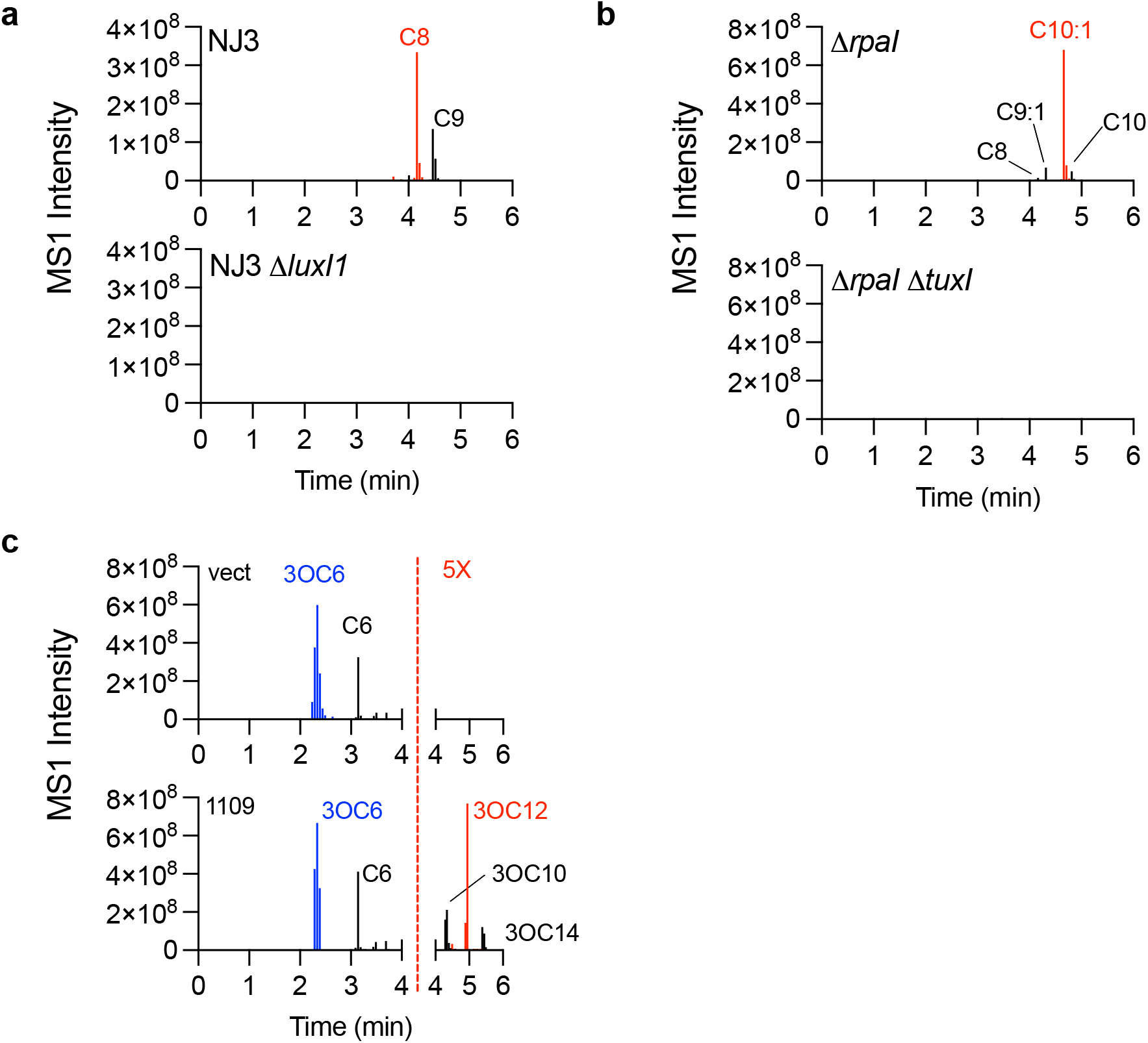
Inventory of AHL compounds associated with cognate LuxI products from select bacteria. Untargeted metabolomics were performed on culture extracts as described in METHODS. **a)** *Mesorhizobium* NJ3 extracts (top panel) have C8-HSL (M+H 228.1593, mass error-0.53 ppm) and minor amounts of C9-HSL (M+H 242.1749, mass error-0.7 ppm). These AHLs are not detected in the strain with a *luxI1* gene deletion (bottom panel). **b)** *R. palustris* TIE-1 Δ*rpaI* (top panel) extracts contain the major AHL C10:1-HSL (M+H 254.1744, mass error-2.64 ppm), in addition to minor amounts of C9:1-HSL (M+H 240.1590, mass error-1.75 ppm), C10-HSL (M+H 256.1901, mass error-2.31 ppm), and C8-HSL (M+H 228.1591, mass error-1.4 ppm). These AHL masses are absent in a strain where the *tuxI* gene is deleted (bottom panel). **c)** *S. odorifera* carrying a vector control plasmid (vect, top panel) produced 3OC6-HSL (M+H 200.1069, mass error-0.6 ppm). However, when expression of its *luxI* homolog SO_1109 was controlled by a constitutive promoter on a plasmid (1109, bottom panel) we could detect 3OC12-HSL (M+H 270.1699, mass error-0.31 ppm) as a major AHL, in addition to lesser amounts of 3OC10-HSL (M+H 298.2006, mass error-2.3 ppm) and 3OC14-HSL (M+H 326.2320, mass error-1.79 ppm). Note that all values to the right of the red dashed line have been multiplied by a factor of 5 to help visualize the results. The extract amount injected for analysis is equivalent to 8 ml of culture in **a)**, and 0.5 ml of culture in **b)** and **c)**.

